# Rice GLUCAN SYNTHASE-LIKE5 promotes Callose deposition in Anthers to maintain proper Male Meiosis Initiation and Progression

**DOI:** 10.1101/2022.05.24.493269

**Authors:** Harsha Somashekar, Manaki Mimura, Katsutoshi Tsuda, Ken-Ichi Nonomura

**Affiliations:** Plant Cytogenetics Laboratory, Department of Gene Function and Phenomics, National Institute of Genetics, Mishima, Shizuoka 411-8540, Japan; Department of Genetics, School of Life Science, The Graduate University of Advanced Studies (SOKENDAI), Mishima, Shizuoka 411-8540, Japan

## Abstract

Callose is a plant cell-wall polysaccharide whose deposition is spatiotemporally regulated in various developmental processes and environmental stress responses. Appearance of callose in premeiotic anthers is a prominent histological hallmark for the onset of meiosis in flowering plants, whose biological role in meiosis is unknown till date. Here we show that rice GLUCAN SYNTHASE LIKE5 (OsGSL5), a callose synthase, localizes on the plasma membrane of pollen mother cells (PMCs), and is responsible for biogenesis of callose in anther locules through premeiotic and meiotic stages. In *osgsl5* mutant anthers mostly lacking callose deposition, aberrant PMCs accompanied by aggregated, unpaired or multivalent chromosomes were frequently observed, and furthermore, a considerable number of mutant PMCs untimely progress into meiosis compared to wild type PMCs. Immunostaining of meiosis-specific protein PAIR2 in premeiotic PMCs revealed precocious meiosis entry in *osgsl5* anthers. The findings of this study bestows new knowledge on function of callose in controlling timing of male meiosis initiation and progression, in addition to roles in microsporogenesis, in flowering plants.

## Introduction

In flowering plants, successful pollen production involves a series of multiple complex steps. An important earlier step is meiosis that takes place within microsporangium or pollen sac, two pairs of which compose an anther in angiosperms (Scott et al. 2004; Zhang and Wilson, 2009). In rice anthers, each of four microsporangia comprises central sporogenous cells (SPCs) surrounded by concentric somatic cell walls four-layered prior to male meiosis, viz. tapetum, middle layer, endothecium and epidermis from inside-out respectively (Fig. 1). After several mitotic divisions, SPCs mature into meiotically competent pollen mother cells (PMCs), which undergo meiosis following DNA replication to halve chromosome number for fertilization, while the tapetal cell (TC) layers provide metabolites and nutrients for neighbouring PMCs and microspores, and eventually ends via programmed cell death (Dickinson and Bell, 1976; Lei and Liu, 2020; Steer, 1977). After anther walls are four-layered, a plant-specific carbohydrate callose fulfils extracellular spaces of a pollen sac chamber or anther locule and surrounds PMCs at premeiotic interphase, which is a prominent histological hallmark for the onset of male meiosis in flowering plants (Shivanna, 2003; Unal et al. 2013).

**Figure 1.**
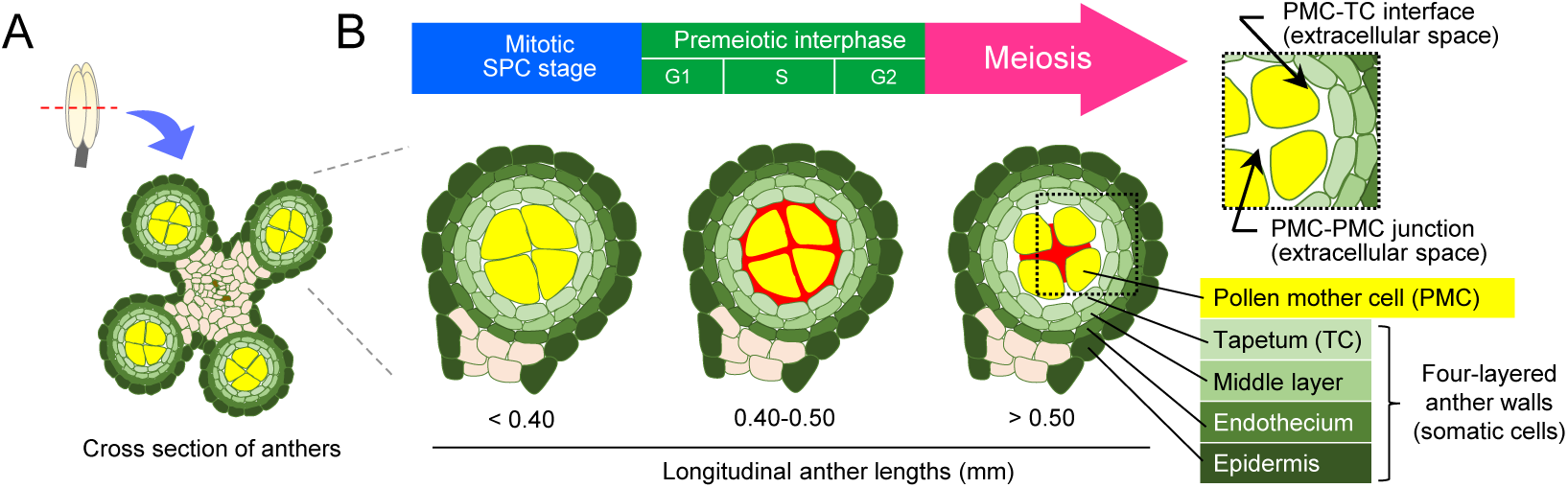
Schematic illustration of anther development and callose accumulation during male meiosis progression. A: Illustration of a cross section of premeiotic anther with four locules. B: Enlarged view of anther locules with constituent cell types, respective anther lengths and their cell cycle status. The red regions in each locule correspond to areas for callose deposition at extracellular spaces observed in this study.

Callose is made up of linear glucose residues of β-1,3 linkages, with some having β-1,6-glucan branches, and functions in various aspects of plant growth and development spatio-temporally (Stone and Clarke, 1992; Chen and Kim, 2009; Zavaliev et al. 2011; Piršelová and Matušíková, 2013; Nedhuka, 2015). For instance, callose is involved in papillae cell-wall materials at bacterial and fungal contact site (Dong et al. 2008; Voigt, 2014) and also secreted at wounded plant tissues (Jacobs et al. 2003), indicating indispensable roles of callose in defence against both biotic and abiotic stresses. In cell-cell signalling, callose regulates the conductivity of plasmodesmata (PD), forming cytoplasmic continuums in plants (Radford et al. 1998; Lucas et al. 2009; Lee and Sieburth, 2010; Zavaliev et al. 2011; Sager and Lee, 2018), and thought to permit selective diffusion of apoplastic signalling (Maltby et al. 1979; Bhalla and Slattery, 1984; Yim and Bradford, 1998). From aspects of plant development, callose is deposited transiently at dividing cell plate during cytokinesis, and aids in primary wall formation between daughter cells (Staehelin and Heplar, 1996; Hong et al. 2001; Thiele et al. 2009). It is also deposited at phloem tissue to control sieve plate development and pore size of sieve tubes (Xie et al. 2011). Towards reproduction, callose helps patterning of pollen aperture and elongation of pollen tubes (Franklin-Tong, 1999; Albert et al. 2011; Qin P et al. 2012; Prieu et al. 2017).

Several reports have revealed the importance of callose accumulation during microsporogensis. In Arabidopsis, *CALLOSE SYNTASE5* (*CalS5*) exerts essential functions during pollen formation stages (Dong et al. 2005). Similarly, rice gene, *GLUCAN SYNTHASE LIKE5* (*OsGSL5*), a homolog of *AtCalS5*, is responsible for pollen growth and development during microsporogensis (Shi et al. 2015). Premature dissolution of callose in the rice *defective callose in meiosis1 (dcm1)* mutant generates abnormal pollen grains with varied size and DNA content as a result of defects in meiotic cytokinesis (Zhang et al. 2018). In rice ovary, OsGSL8 symplasmically controls unloading carbohydrates into pericarp cells of developing ovary, in addition to regulating vascular cell patterning (Song et al. 2016). In Arabidopsis, the amount of phloem-mobile GFPs, able to be unloaded onto all gynoecium cells before female meiosis, are extremely reduced around tetrad spores, suggesting physical isolation of meiocytes from other gynoecium cells probably by callose accumulation (Werner et al. 2011). In contrast, little is known about the roles of callose accumulation in premeiotic anther development and male meiosis, despite of its noticeable amounts and cross-species conservation in land plant anthers (Musial and Koscinska-Pajak, 2017; Sager and Lee 2018; Seale, 2020).

Our group previously reported that MEIOSIS ARRESTED AT LEPTOTENE2 (MEL2), an RNA recognition motif protein, functions in timely transition of spore mother cells to the meiotic cycle in rice (Nonomura et al. 2011). Interestingly, one of significantly downregulated genes in premeiotic *mel2* mutant anthers was *OsGSL5*, detected by transcriptome (Mimura et al. 2021) and reverse transcription quantitative PCR (RT-qPCR) of this study (Supporting table 1), and as expected, callose accumulation was largely eliminated from *mel2* anther locules during premeiosis and meiosis (Fig. S2). Among 10 rice *GSL* genes, *OsGSL5* is an only gene expressing preferentially and abundantly in anthers during meiosis and post-meiosis (Yamaguchi et al. 2006; Shi et al. 2015) (Fig. S3). These findings suggest an unknown association of callose deposition with meiotic entry control in plants.

This study demonstrates that *OsGSL5* is responsible for hyper callose accumulation in extracellular spaces of anther locules at premeiosis and early meiosis, in addition to the role in late meiosis and pollen development as previously reported (Shi et al. 2015), and has an indispensable role in proper initiation of male meiosis in rice.

## Results

### OsGSL5 impacts on pollen viability and seed fertility

The *osgsl5* mutation is reported to cause male gametophytic lethality (Shi et al. 2015), leading to no homozygous plants from self-pollination of heterozygous plants. Thus, to assess OsGSL5 impact on male sporogenesis and meiosis, we exploited CRISPR-Cas9 strategy to directly induce biallelic mutations in *OsGSL5* locus. Out of 32 CRISPR-edited plantlets screened, we obtained two independent knocked-out lines, *osgsl5-2* and *osgsl5-3*, in which 1bp (A) was inserted on the 25th exon and 2bp (TA) were deleted on the 14th exon, respectively (Fig. 2A). Both *osgsl5* lines were largely comparable to wild type (WT) plants in vegetative growth and panicles and spikelets morphologies under the same condition (Fig. 2B, C), except for two weeks-later heading. However, both lines set no seeds, while WT plants had 63% fertility (Fig. 2C Table S2). Pollen viability was 0.5% and 1.2% in *osgsl5-2* and *osgsl5-3*, respectively, whereas it was 90.4% in WT (Figs. 2D, S5). These results were largely consistent to those previously reported (Shi et al. 2015).

**Figure 2.**
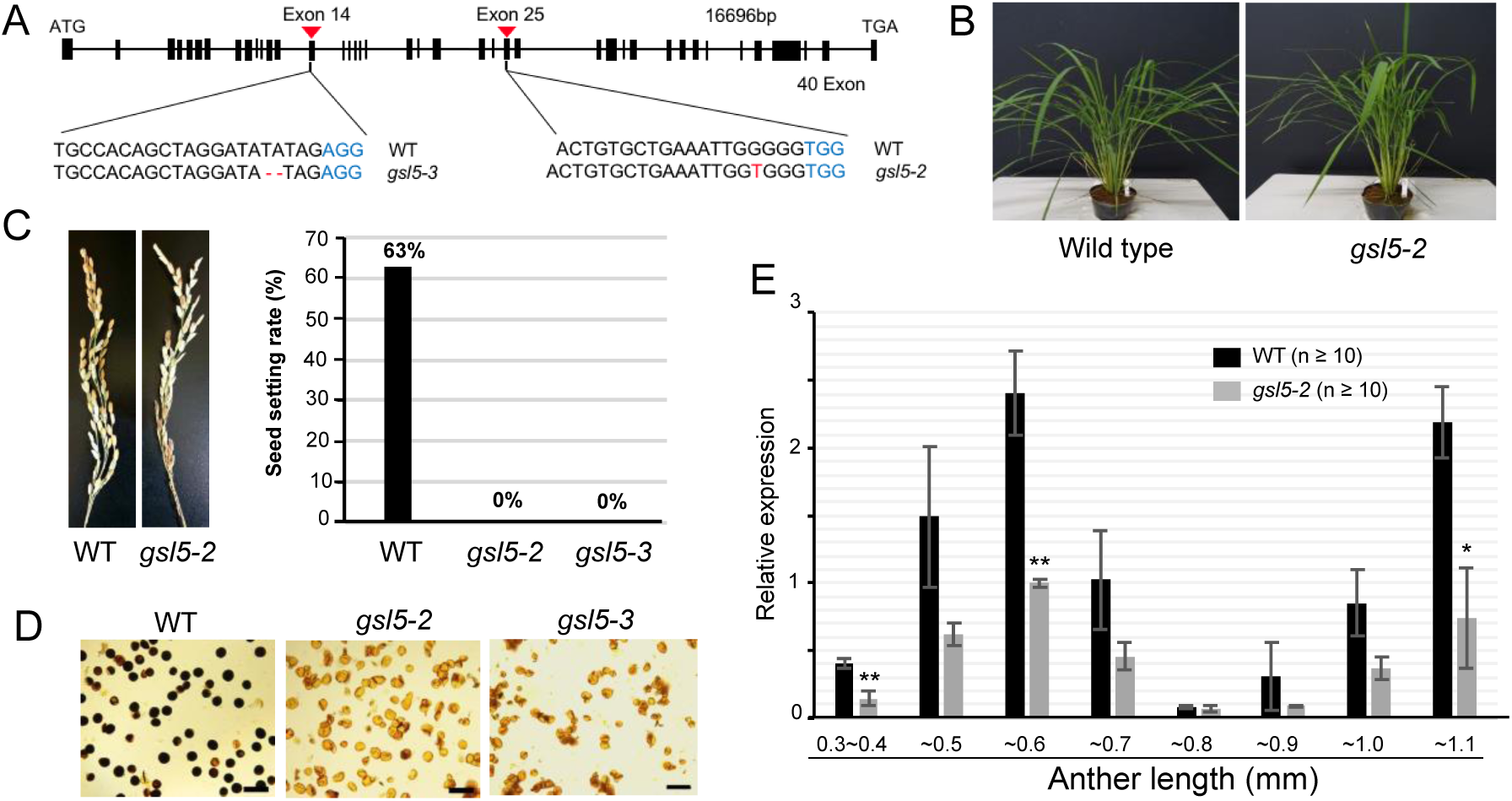
The *osgsl5* mutant phenotype and *OsGSL5* gene expression. A: Nucleotides deletion and insertion sites in *osgsl5-2* and *osgsl5-3* knock-out mutants. B: The vegetative plant growth was comparable between wild type and *osgsl5-2*. C: Images of panicles at seed filling stage of wild type and *osgsl5-2* (left), and the graph showing seed setting rate of wild type, *osgsl5-2* and *osgsl5-3* (right). D: Pollen viability test by KI_2_ staining of WT (left), *osgsl5-2* (middle) and *osgsl5-3* (right). Darkly and faintly stained grains were categorized as viable and nonviable pollen, respectively. Bar=40μm. E: The relative expression levels of *OsGSL5* transcripts in WT and *osgsl5-2* anthers by RT-qPCR. Errors bars each indicate the standard deviation of the mean of 3 biological replicates. The n value in parentheses is the number of florets used in each replicate. Asterisks indicate significant differences (P<0.05) between WT and *osgsl5-2* (student’s t-test).

In WT plants, the *OsGSL5* transcript level was highest in anthers, while undetected in vegetative organs and slightly detectable in pistils (Fig. S6). It was significantly enhanced in 0.4-0.5mm anthers at premeiotic interphase, three-fold more than the level in younger 0.3-0.4mm anthers which contain SPCs proliferating mitotically (mitotic SPC stage). The expression peaked at 0.5-0.6mm anthers around leptotene to pachytene. In 0.6-0.7mm anthers around diplotene to telophases I, the *OsGSL5* level was reduced to half or less of that at the former stage, and again elevated in 0.8-1.1mm anthers at tetrad and microspore stages (Fig. 2E). Though the *osgsl5-2* mutant developed anthers with normal appearance (Fig. S7), the *OsGSL5* transcript level was lowered in all *gsl5-2* anthers examined, and significantly reduced at premeiotic interphase, early prophase I and microspore stages (Fig. 2E).

Considering downregulation of *OsGSL5* transcription and reduced fertilities of pollen and seeds similarly in two independent lines, we concluded that CRISPR-Cas9-mediated frameshift mutations within *OsGSL5* coding sequence caused all above phenotypes.

The anther length is broadly utilized as a standard to assess meiotic events in many angiosperm species including rice, as it has a rough collinearity with respective meiotic stages (Itoh et al. 2005) (Fig. 1). To confirm that *osgsl5* mutant anthers retain this collinearity, we examined the expression profiles of tapetum-specific genes, *TIP2* (Fu et al. 2014) and *EAT1* (Niu et al. 2013). The expression peaks of both genes appeared at a similar level in both WT and *osgsl5-2* anthers (Fig. S7A), noteworthily in which bimodal *EAT1* expression peaks at both early meiosis and tetrad stages previously reported (Ono et al. 2018) were completely maintained even in *osgsl5-2* anthers. Furthermore, no difference in the layered structure of premeiotic anthers was observed between WT and *osgsl5* plants (Fig. S7B). These results indicate that the *osgsl5* mutation unlikely affects the collinearity between anther lengths and meiotic stages, in addition to earlier anther morphogenesis. Thus we utilized the anther length as a standard for comparison of meiotic events between WT and *osgsl5* mutants below.

### OsGSL5 impacts on callose accumulation in premeiotic and meiotic anther locules

We monitored the detailed pattern of callose accumulation in rice meiotic anthers by aniline blue that specifically stains β-1,3-glucan chains. The callose amount in WT anther locules, which was at an undetectable level during mitotic SPC stage (Fig. 3A), started to fulfill the extracellular spaces between cell walls and cell membranes at both PMC-PMC junctions and PMC-TC interfaces (Fig. 3B). When PMCs entered into meiotic leptotene and zygotene, callose deposition was limited to PMC-PMC junctions (Fig. 3C, D). At subsequent pachytene to diakinesis stages, it can be seen enclosing PMCs with rounder shape (Fig. 3E-G). At dyad and tetrad stages, callose was detected on newly formed equatorial cell plates in addition to outer cell surfaces (Fig. 3H-I). After release of microspores to anther locules, callose can be seen fully enclosing microspores (Fig. 3J). In both *gsl5-2* and *gsl5-3* anthers, callose signals were extremely diminished through all above stages (Fig. 3K-T).

**Figure 3.**
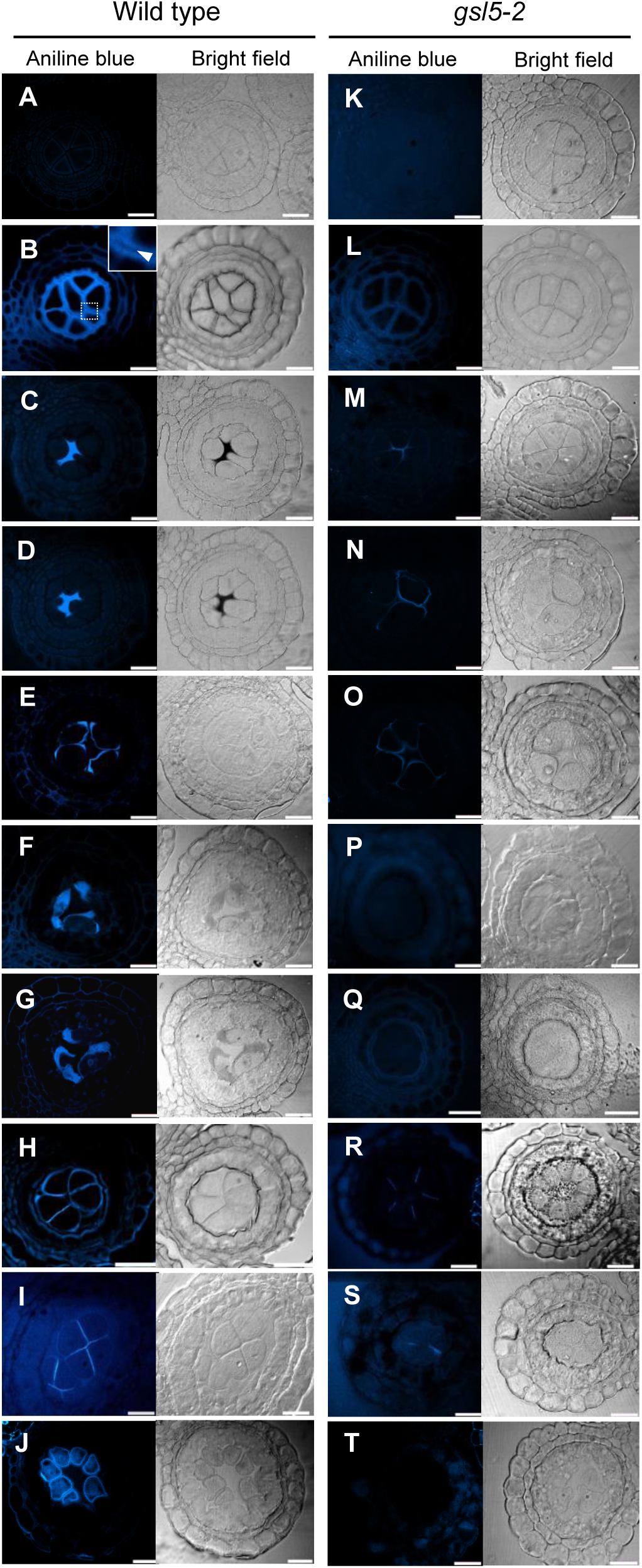
Callose accumulation during male meiosis in WT and *osgsl5-2* anthers. Callose accumulation pattern was monitored by aniline blue staining at different male meiosis substages. A and K: Mitotic SPC stage, B and L: Premeiotic interphase, C and M: Leptotene, D and N: Zygotene, E and O: Pachytene, F and P: Diplotene, G and Q: Diakinesis, H and R: Dyad, I and S: Tetrad, J and T: Microspore. An arrowhead in the inset in B indicates the cell wall unstained with aniline blue. Bar=20μm.

A beginning stage of callose accumulation was further observed with the anti-β-1,3-glucan (callose)-directed antibody, to trace differences in callose amount more sensitively. In young 0.4-mm or less anthers, four microsporangia of a same anther sometimes show different callose patterns with each other (Fig. S8), suggesting it being at the very beginning of premeiotic callose accumulation. In this stage, locules frequently retained callose limited to PMC-PMC junctions (Fig. S8), implicating that premeiotic callose accumulation initiates around PMC-PMC junctions, but not at PMC-TC interfaces.

### Localization of Callose and OsGSL5 protein in anther locules

To investigate spatio-temporal localization of OsGSL5 protein, we produced anti-OsGSL5 antiserum (Fig. S4) to perform co-immunostaining with the anti-callose antibody on anther sections. After initiation of callose accumulation at PMC-PMC junctions (Fig. S8), callose fulfilled extracellular spaces of both PMC-PMC and PMC-TC regions in anther locules (Fig. 4A), as observed in aniline blue staining (Fig. 3B). In contrast to callose staining, OsGSL5 localization was limited only at PMC-PMC junctions in the same anther section (Fig. 4A). In subsequent leptotene, the thick callose signal declined again at PMC-TC interfaces and became almost overlapped with OsGSL5 localization at PMC-PMC junctions (Fig. 4B-C). Through all stages observed, the strongest linear OsGSL5 signals were always observed at optically sectioned edges of PMCs (open arrowheads in Fig. 4), suggesting their association with the plasma membrane, well consistent to the fact that OsGSL5 and its orthologs are membrane-anchored proteins containing 15 transmembrane domains (Yamaguchi et al. 2004; Shi et al. 2015) (Fig. S4A). This trend became more obvious at subsequent pachytene-diakinesis stages, where PMCs gradually took a spherical shape (Fig. 4D-E). During these stages, OsGSL5 localization (Fig. 4D, E) was gradually corresponding to callose deposition on PMC surfaces (Figs. 3E-F, 4D-E). The OsGSL5 signal was at undetectable level in *osgsl5* mutant anthers (Fig. S9).

**Figure 4.**
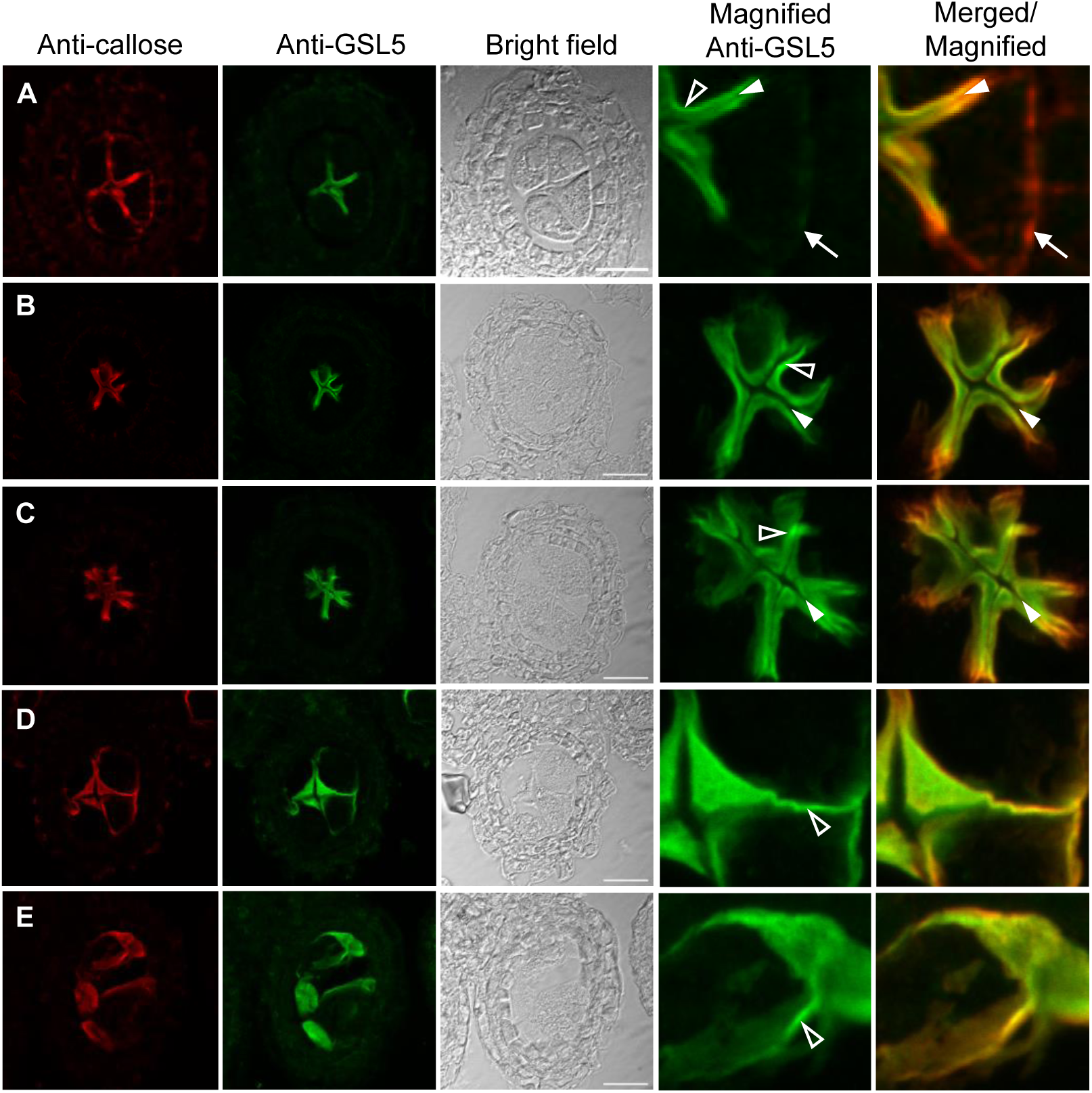
Co-immunostaining of callose and OsGSL5 protein in WT anthers. A: Premeiotic interphase, B: Leptotene, C: Zygotene, D: Pachytene, E: Diplotene-Diakinesis. Bar=20μm. In panel A, white arrows show the region retaining callose deposition (red), but lacking OsGSL5 localization (green), at PMC-TC interface. The closed and open arrowheads indicate unstained cell-wall regions and linearly aligned OsGSL5 signals on PMC plasma membranes, respectively.

These results strongly support that OsGSL5 takes part in callose biosynthesis during meiosis stages, and taken together with RT-qPCR results, reconfirm that the *osgsl5* mutations are null alleles.

### OsGSL5 impacts on male meiosis progression and chromosome behaviours

The chromosome behaviour was assessed at respective meiotic stages determined by both anther lengths and chromosomal morphologies on plastic-embedded sections. In WT, PMCs began meiotic chromosome condensation and displayed thin thread-like appearance by leptotene (Fig. 5A-C). Chromosomes were further condensed during zygotene (Fig. 5D), at which homologous chromosomes begin to be synapsed. At pachytene when synapsis is completed, PMCs displayed thick-threads of homologous pairs (Fig. 5E, E’). Bivalent chromosomes were further condensed through diplotene and diakinesis (Fig. 5F, G), and either of homologous pair was delivered to opposite poles during meta/anaphase I, and eventually to either cell of the dyad (Fig. 5H). In both *gsl5-2* and *gsl5-3* anthers, a conspicuous abnormality on male meiotic chromosomes emerged during prophase I (Fig. 5I-O), in which two different types of PMCs were contained together within a same anther - one type carrying meiotic chromosomes with seemingly normal appearance, named wildtype-like PMC (wl-PMC) (Fig. 5L, M, M’_wl), and another displaying aberrant behaviors of meiotic chromosomes, such as tight aggregations and impaired homologous pairings, named aberrant PMC (ab-PMC) (Fig. 5L-O, M’_ab). The chromosome spreading method further enabled to clarify the abnormality in *osgsl5* ab-PMCs (Fig. S10).

**Figure 5.**
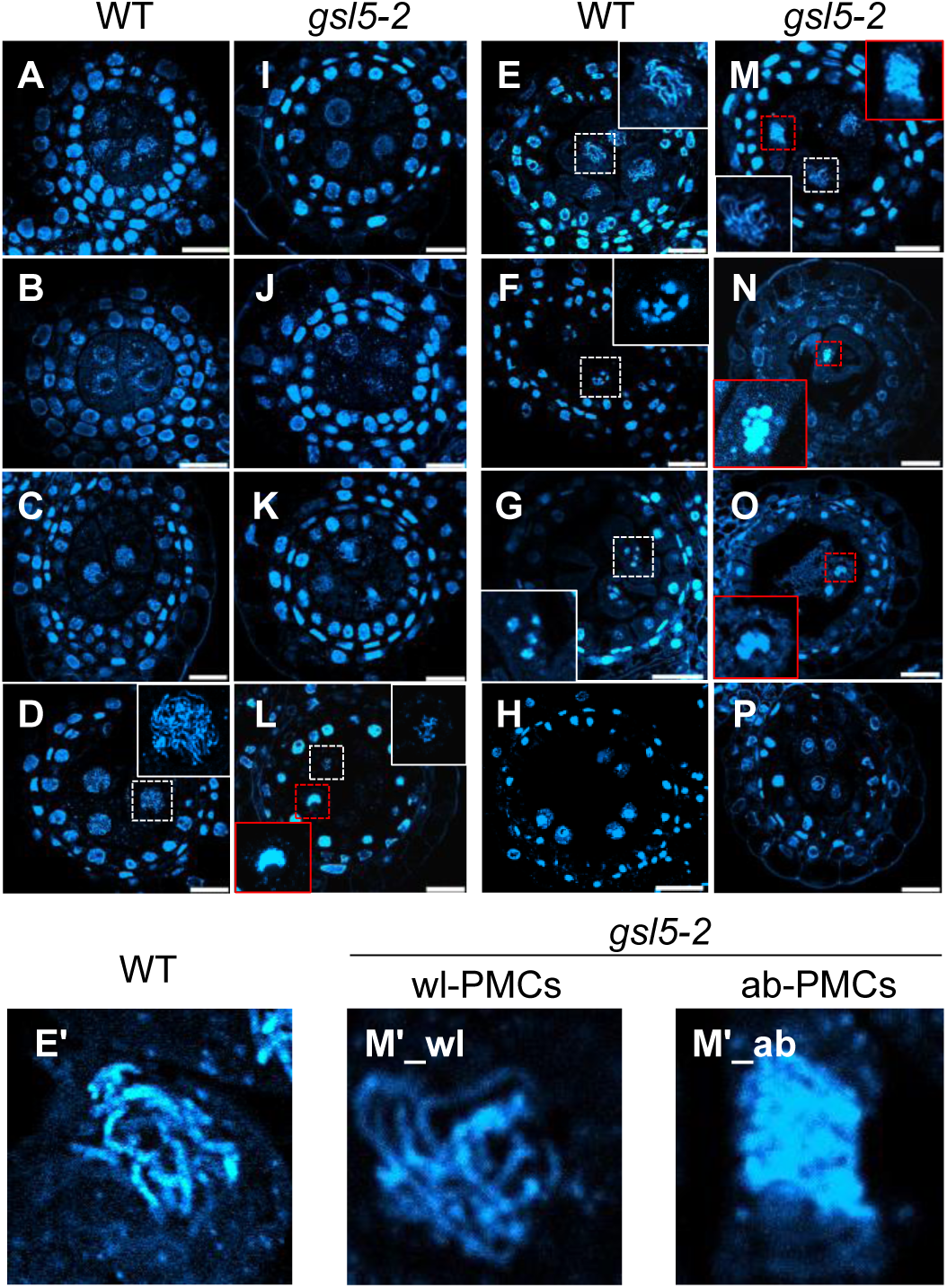
Dynamics of chromosomal behaviour in WT and *osgsl5-2* anthers. Plastic-embedded anthers cross sections stained with DAPI for chromosomes observation. A and I: Mitosis, B and J: Interphase, C and K: Leptotene, D and L: Zygotene, E and M: Pachytene, F and N: Diplotene, G and O: Diakinesis, H and P: Dyad. Bar=20μm. Note that *osgsl5-*2 mutant anthers contained two types PMCs with respect to chromosome appearance (wl-PMC and ab-PMC, see the text). The insets in each panel are magnified views of nuclei enclosed with dashed squares, in which white and red squares indicate nuclei of wl-PMCs and ab-PMCs, respectively. The insets in panels E and M are further magnified and shown in E’, M’1 and M’2, as examples of WT PMC, wl-PMC and ab-PMC.

In above observations, we noticed the *osgsl5* mutation somewhat affected time-course progression of male meiosis, in addition to segregation of two PMC types. Thus, to make the *osgsl5* impact on meiosis progression clearer, the frequency of each meiosis stage observed in PMCs was plotted along anther lengths. In 0.4-to 0.8-mm *osgsl5* anthers, ab-PMCs appeared irregularly in range of 8.2-46.2% of all PMCs observed (Fig. 6A, Table S3). Another point of interest was a precocious initiation of several meiotic stages in not all, but a part of wl-PMCs, compared to WT PMCs. Interestingly, despite a subset of wl-PMCs displaying chromosomal features particularly of zygotene to dyad were observed earlier than WT PMC stages (asterisks in Fig. 6A), male meiosis was completed similarly in 0.9mm anthers of both WT and *osgsl5* plants (Fig. 6A, Table S3).

**Figure 6.**
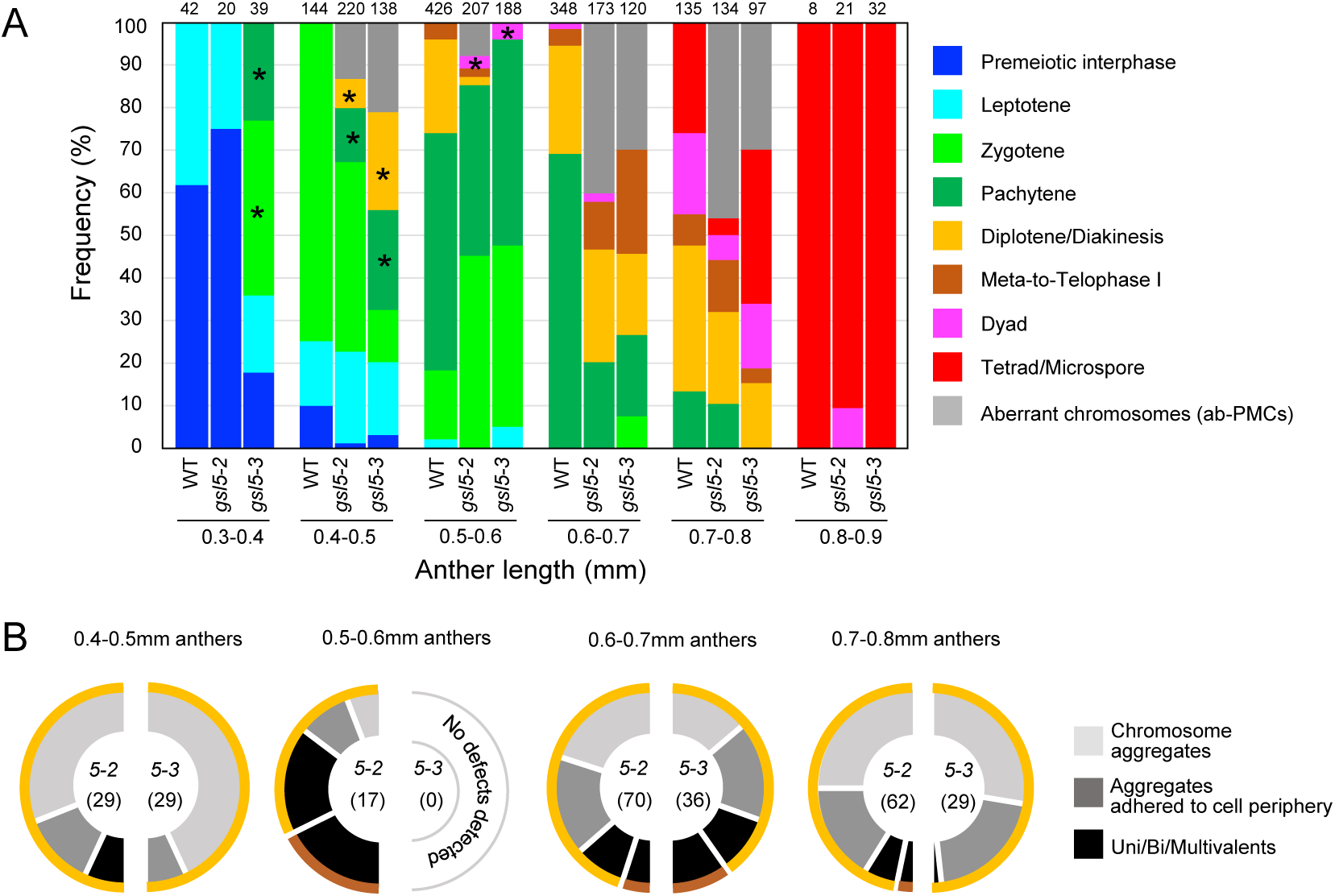
Male meiosis progression and ab-PMC appearance in WT and *osgsl5* anthers. A: Frequency of PMC stages observed with respect to anther lengths in WT and *osgsl5* mutants, in which the frequency of ab-PMCs in *osgsl5* anthers was shown as gray bars. The number at the top of each bar indicates the absolute number of cells counted. The wl-PMC stages marked with asterisks indicate that those stages emerged much earlier than comparable stages observed in WT anthers. B: Each half-donut graph indicates frequency of three different classes for aberrant chromosomal morphologies and behaviors in *osgsl5-2* or *osgsl5-3* ab-PMCs observed along respective anther lengths. Colored outer rims on a half donut indicate meiotic stages of wl-PMCs that concomitantly emerge with respective classes of ab-PMCs. Definition of outer rim colors is consistent to that in A. The number in parentheses represents the absolute number of cells counted.

In 0.4-0.5mm *gsl5* anthers, >85% of ab-PMCs retained chromosomal aggregates, and concomitantly appeared with diplotene/diakinesis-like wl-PMCs (Figs. 6B, S10, Table S3). In 0.6-0.7mm *gsl5* anthers, about >27% of ab-PMCs had more condensed univalents and/or multivalents in addition to normal bivalents, concomitant with wl-PMCs retaining diplotene/diakinesis-or meta/telophase I-like chromosomes (Figs. 6B, S10, Table S3). In 0.7-0.8mm anthers, >80% ab-PMCs again displayed less-condensed aggregated chromosomes concomitantly with diplotene/diakinesis-like wl-PMCs (Figs. 6B, S10, Table S3). The reason was ambiguous, but it may suggest that aberrant aggregations of chromosomes at early prophase I resulted in uni/multivalent formation at later stages, and that ab-PMCs retaining uni/multivalents were abortive and undetected during late prophase I.

### *osgsl5* mutation caused precocious initiation of male meiosis

An earlier occurrence of meiotic prophase-I stages frequent in *osgsl5* anthers (Fig. 6) raises a possibility that it is attributable to defects in premeiotic events. To test this hypothesis, we performed PAIR2 immunostaining. Rice PAIR2 promotes homologous chromosome synapsis, and its accumulation within the nucleus is reported to initiate during premeiotic interphase, just as following the initiation of premeiotic DNA replication (Nonomura et al. 2006). Furthermore, the transcriptional level of *PAIR2* gene was unaffected by *osgsl5* mutation through all meiotic stages (Fig. S11). Thus, PAIR2 can be used as a maker to infer the timing of replication initiation in anthers at premeiotic interphase.

PAIR2 nuclear signals were classified into three classes by their intensity; absent (class I), faint (class II) and strong (class III) (Fig. 7A), of which the class II was supposed to be a stage following premeiotic replication initiation. In WT 0.30-0.45mm anthers observed, 80% of PMCs showed no PAIR2 signal (class I), suggesting the cells being at mitotic SPC stage or before replication, and only 20% showed class II signals. In contrast, in anthers with the same lengths, around 60-70% of *osgsl5* PMCs showed either class II or III signals (Fig. 7B, C). These results clearly indicate that OsGSL5 has an impact on timely initiation of premeiotic events, such as PAIR2 loading that contemporises with DNA synthesis during premeiotic interphase.

**Figure 7.**
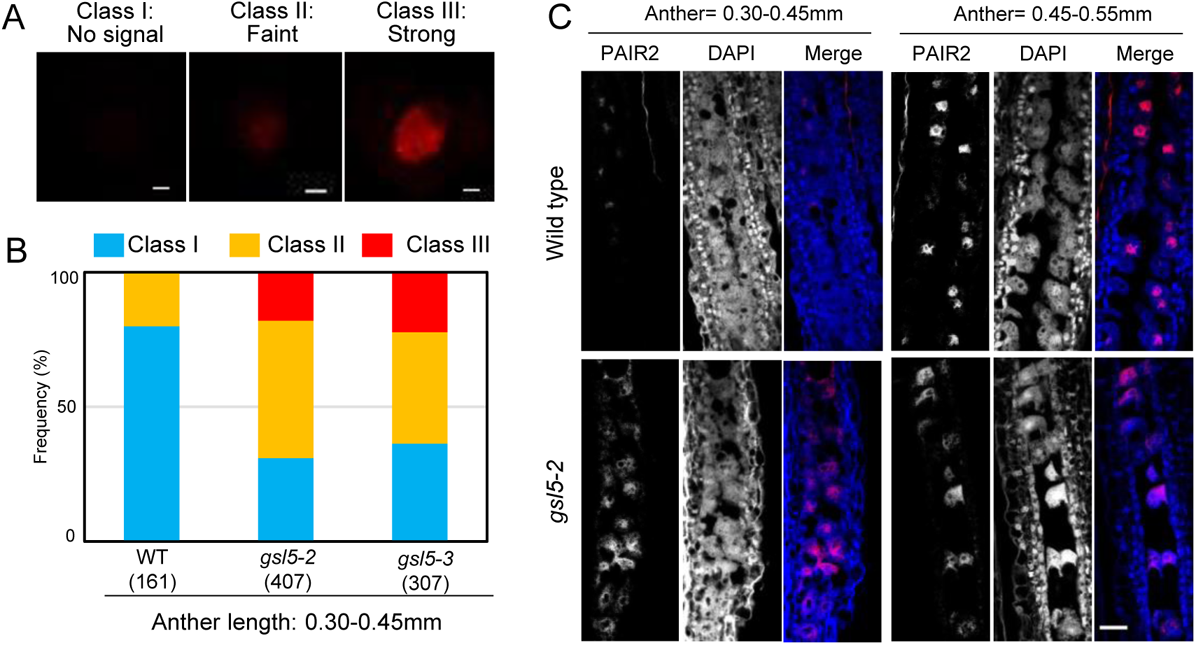
Precocious accumulation of PAIR2 in *osgsl5* PMC nuclei. A: Immunostaining of PAIR2 in isolated premeiotic PMCs. The degree of PAIR2 accumulation in the PMC nucleus was categorized into three classes based on the immunofluorescent intensity. Bar=2.5μm. B: The rate of three PAIR2 classes in each of WT, *osgsl5-2* and *osgsl5-3* premeiotic anthers. C: Immunostaining of PAIR2 on longitudinal premeiotic anther sections with anti-PAIR2 antibody. Bar= 50μm.

### *osgsl5* mutation disrupts homologous synapsis

Next, we asked whether key meiotic events such as homologous chromosome synapsis was affected in *osgsl5* PMCs or not. PAIR2 was normally loaded on meiotic chromosomes normally in both WT PMCs (n=168) and *osgsl5-2* PMCs observed (n=134) (Fig. 8A), which further ensured that premeiotic accumulation of PAIR2 in PMC nuclei occurs normally even in *gsl5* mutants (Fig. 7). In contrast, loading of ZEP1, which is a transverse filament component of synaptonemal complex and governs meiotic crossover numbers in rice (Wang et al. 2010), was severely diminished in all *osgsl5-2* PMCs at zygotene and pachytene (n=85), while it was constantly observed in all WT PMCs in same stages (n=92) (Fig. 8B) and the *ZEP1* gene expression was comparable between WT and *osgsl5-2* anthers (Fig. S11). Though a chromosomal aggregate characteristic of ab-PMCs was hard to be distinguished at these stages, the result suggests that failed ZEP1 loading took place in both wl- and ab-PMCs in *osgsl5* anthers.

**Figure 8.**
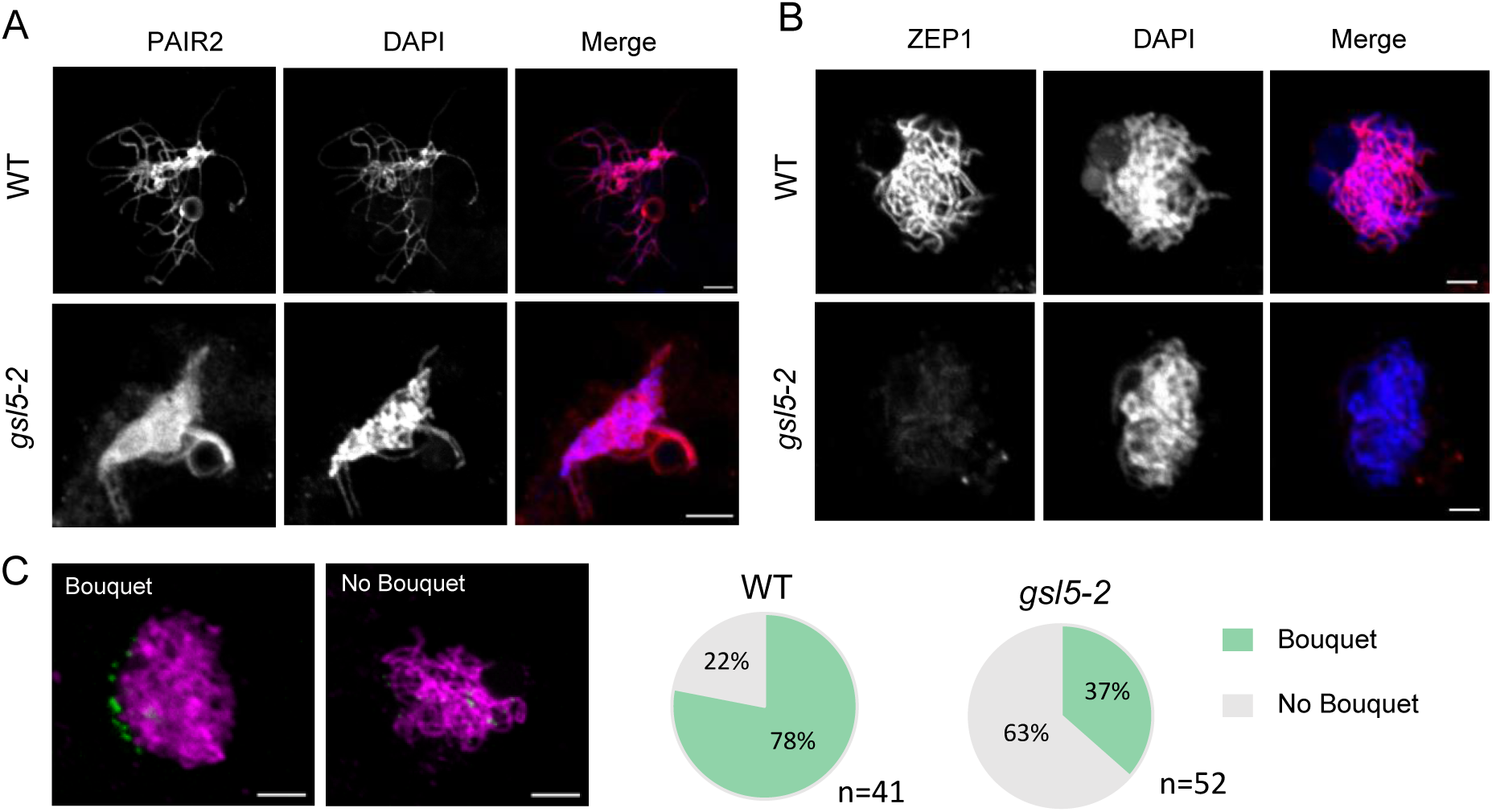
ZEP1 loading to and bouquet frequency of meiotic chromosomes affected, but PAIR2 loading unaffected in *osgsl5* anthers. A: Immunostaining of ZEP1 in WT (n=92) and *ososgsl5-2* (n=92). ZEP1 loading onto chromosomes was inhibited in *osgsl5* mutant. B: Immunostaining of PAIR2 in WT (n=168) and *osgsl5-2* (n=134). PAIR2 loading is unaffected in *osgsl5* mutant. C: Bouquet structures visualized by immunostaining of telomere specific OsPOT1 during leptotene-zygotene transition (0.50-0.55mm anther) in PMCs of WT (n=41) and *osgsl5-2* (n=52). Bar=5μm.

Telomere bouquet is a chromosomal arrangement important for meiotic homologous pairing and synapsis in many eukaryotes including rice (Zhang et al. 2017). In anthers around leptotene and zygotene, only 37% PMCs displayed the bouquet in *osgsl5-2* mutant (n=52), while 78% did in WT (n=41) (Fig. 8C). In 0.60-0.70mm anthers, no bouquet was observed both in WT and *osgsl5-2* mutant (Fig. S12), suggesting the *osgsl5* mutation restricted bouquet formation, but not dissolution.

### Altered transcript levels of key meiotic and callose metabolizing genes in *gsl5* anthers

The transcriptional levels of 13 genes having key roles in meiosis and two genes involved in callose metabolism were quantified in meiotic anthers by RT-qPCR. Of 13 genes, *PAIR3* and *MER3* were significantly upregulated at early meiosis (0.3-0.5mm anthers), and *REC8* was upregulated significantly at mid and late meiotic stages (0.6-1.1mm anthers) in *osgsl5-2* anthers (Fig. S11). The transcript levels of other 10 meiotic genes were comparable between WT and *osgsl*5 (Fig S11).

Noteworthily, two callose metabolizing genes, *Osg1* gene encoding a tapetum-specific *β*- 1,3-glucanase (Wan et al. 2011) and *UGP1* gene encoding an UDP-glucose phosphorylase involved in biosynthesis of cell wall components including callose (Chen et al. 2007), were both downregulated in *osgsl5-2* anthers through all meiotic stages (Fig. 9).

**Figure 9.**
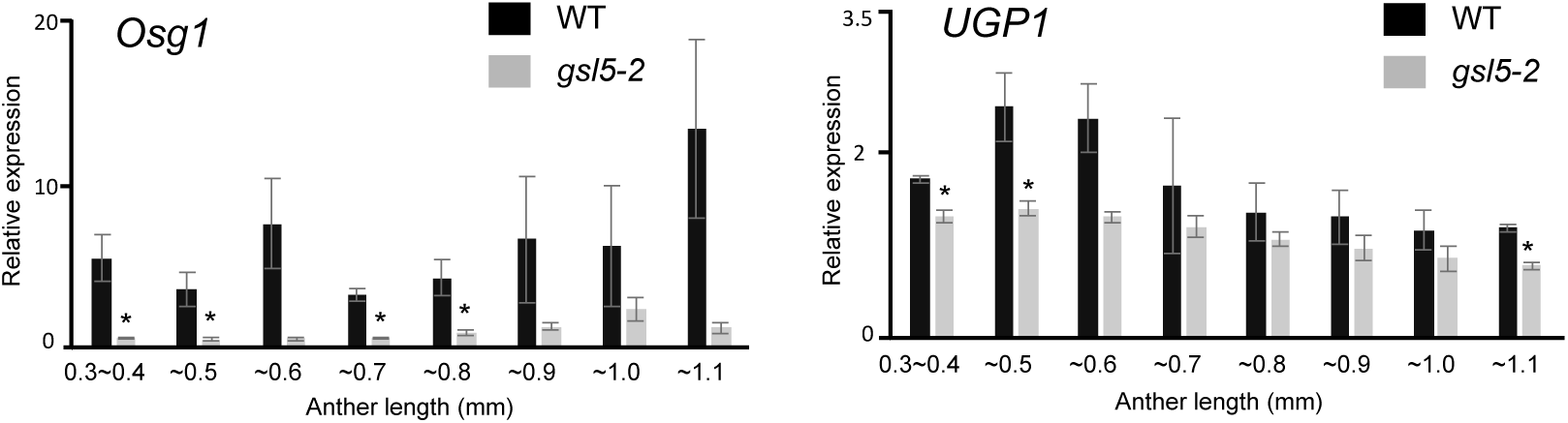
Reduced expression of rice *Osg1* and *UGP1* genes in *osgsl5-2* anthers. Relative expression levels of callose metabolizing genes, *Osg1* and *UGP1*, in WT and *osgsl5-2* anthers by RT-qPCR. Errors bars indicate standard deviations of means of three biological replicates. Asterisks indicate significant differences (P<0.05) between WT and *osgsl5-2* (student’s t-test).

## Discussion

### OsGSL5-dependent callose accumulation in rice anthers during meiosis

This study gained important insights into OsGSL5 callose synthase responsible for callose deposition at extracellular spaces of anther locules during premeiotic and meiotic stages (Fig. 3). Double immunostaining of OsGSL5 and callose revealed that subcellular OsGSL5 localizations are restricted on PMC plasma membrane facing to PMC-PMC junction, where multiple PMCs meet with each other along the central axis of anther locules, and served callose polysaccharides to extracellular spaces of PMC-PMC junctions through premeiotic interphase and early prophase I (Fig. 4A-D). Another important point is that during premeiosis, callose deposition was observed at PMC-TC interfaces in addition to PMC-PMC junctions, whereas OsGSL5 localization was limited to PMC-PMC (Fig. 4A). Given that callose accumulation begins at PMC-PMC junctions, but not at PMC-TC interphases (Fig. S8), callose synthesized at PMC-PMC junction is likely supplied for fulfilling PMC-TC interfaces during premeioptic interphase, further affirming PMC-PMC junctions as a callose-producing center.

Fluctuation in callose levels, as above mentioned, generally involves two counteracting enzymatic activities: β-1,3-glucan synthases and hydrolases (Frankel et al. 1969; Stieglitz and Stern, 1973). A rice β-1,3-glucanase, Osg1, functions in timely callose degradation on pollen grains and impact on pollen fertility (Yamaguchi et al. 2002; Wan et al. 2011), and *Osg1* gene expression was reduced in *osgsl5* mutant (Fig. 9), likely suggesting a positive feedback regulation of *Osg1* transcription by elevated callose levels to maintain callose homeostasis in anthers. Rice *UGP1* is a UDP-glucose pyrophosphorylase that catalyzes production of UDP-glucose, a substrate of glucan synthases, including GSL5. In *UGP1* RNAi plants, callose accumulation in anther was significantly diminished during both meiotic and post-meiotic stages (Chen et al. 2007) which is consistent with our results (Fig. 3). Thus, it is possible that UDP-glucose catalyzed by *UGP1* is used for *GSL5*-dependent callose synthesis, leading to callose accumulation during meiosis and post-meiosis, and *Osg1* probably acts in callose fluctuation antagonistically to the UGP1-GSL5 pathway.

### Impact of callose accumulation on male meiosis initiation

A key insight gained with this study is the impact of OsGSL5 on male meiosis initiation and progression (Figs. 6, 7). In addition to defects in meiosis time-course, *osgsl5* mutant included ab-PMCs that exhibited several other defects in chromosome behaviour and condensation, homologous synapses and reduced bouquet structures, along with wl-PMCs that exhibited a normal appearance of meiotic chromosomes (Figs. 5, 8, S10). Even if a part of *osgsl5* PMCs passed through meiosis, callose deposition is defective also on all surviving tetrad spores (Fig. 3T), resulting in male sterility as previously reported (Shi et al. 2015).

Shi et al. (2015) concluded that OsGSL5 is responsible for callose accumulation at post meiosis stages, but not during early meiosis. However, this conclusion was led only by a snapshot of whole-mount staining of anthers with aniline blue, but not of sectioning, perhaps resulting in an oversight of OsGSL5 impact during premeiotic interphase and prophase I stages. Similarly, a frequent appearance of survival spores, probably derived from wl-PMCs, may explain the reason why the impact of callose function during male meiosis was underestimated in previous studies using mutant plants lacking callose synthesis.

Microscopic observations of *osgsl5* mutant PMCs implicated that wl-PMCs with meiotic chromosomes lacked ZEP1 loading (Fig. 8B), but achieved seemingly normal bivalent formation and disjunction of homologous pairs in meiosis I (Fig. S10F, I). It is not surprised, because in rice *zep1* mutants, bivalents were formed at the normal level and a considerable number of tetrads were formed, while the viability of gametes significantly reduced probably due to aberrant chromosomal condensation in microspores (Wang et al. 2010). Rather, it is a wonder why meiotic chromosomes of *osgsl5* wl-PMCs loose the capacity for ZEP1 loading. Furthermore, frequent appearance of ab-PMCs is difficult to be explained only by the loss of ZEP1, because of no appearance of such aggregates reported in *zep1* mutants (Wang et al. 2010). A similar loss of ZEP1 loading is observed in *mel2* mutant PMCs (Nonomura et al. 2011), in which *OsGSL5* expression is significantly reduced (Fig. S1). Furthermore, PMCs at various cell cycle stages segregated in a same *mel2* anther, due to asynchronous initiation of DNA replication (Nonomura et al. 2011). This observation may account for appearance of two different PMC types in *gsl5* anthers (Figs. 5, 6), taking place together with precocious male-meiosis entry (Fig. 7). Rice LEPTOTENE1 (LEPTO1), a type-B response regulator that participates in establishing key features of meiotic leptotene chromosomes. In *lepto1* mutant anthers, expression levels of both *OsGSL5* and *UGP1* genes were reduced significantly and callose depleted during meiosis (Zhao et al. 2018). Interestingly, the loading of important meiotic chromosome elements, such as OsREC8, OsAM1 and ZEP1, was also defected in *lepto1* (Zhao et al. 2018). The past observations and findings of this study together strongly implicate that callose filling the extracellular spaces of anther locules is an important step for proper PMC differentiation and/or male meiosis initiation, while the underlying mechanisms have been remained elusive.

Recent studies often suggest the role of callose accumulation in male meiosis via cross-talking among anther locular cells (Plackett et al. 2014; Zhai et al. 2015; Liu et al. 2017; Huang et al. 2019; Lei and Liu 2020). PMCs are interconnected with each other and with surrounding TCs through PD or cytomictic channels at early meiosis (Heslop-Harrison, 1964; Mamun et al. 2005; Mursalimov et al. 2010; Mursalimov et al. 2013). However, during transition to meiosis, such intercellular connections are solved/blocked probably by hyper callose accumulation (Sager and Lee, 2018), probably by hyper accumulation of callose in anther locules. In addition to controlling symplastic pathway, callose accumulation is thought to function as a molecular filter for signalling from TCs to PMCs via apoplastic pathway (Clement & Audran, 1995; Roschzttardtz et al. 2013. Biochemical evidence further suggest that callose deposition can alter permeability and plasticity of cell walls in coexistence with cell-wall components like cellulose (Abou-Saleh et al. 2018). The above findings imply that hyper callose accumulation has a potential to bring dramatic microenvironmental changes to its surrounding PMCs via controlling symplastic or apoplastic pathways or both, which entails to be confirmed in future studies.

In summary, this study demonstrates that *GSL5*-dependent callose deposition during meiosis is crucial for proper timing of meiosis initiation and subsequent progression, and upon callose depletion at this point of time perturbs normal meiosis onset and consequently pose severe impact on several meiotic events. This study sheds light on the importance of callose in meiosis of higher plants which is an important progress in the field of plant reproductive biology, and is a first step towards understanding the mechanistic basis of *GSL5* and callose function in meiosis initiation.

## Materials and Methods

### Plant materials and growth conditions

For target mutagenesis of *OsGSL5* gene (*Os06g0182300*), potential CRISPR guide-RNA sequences were designed using CRISPR-P v2.0 software (Liu et al. 2017). Double stranded DNAs were produced from oligo DNA pairs of gRNAF1/gRNAR1 and gRNAF2/gRNAR2 for *osgsl5-2* and *osgsl5-3* (Table S2), respectively, by annealing. After cloning into pU6 vector, the *pU6* promoter-fused double-stranded DNA was transferred to pZD shuttle vector (Mikami et al. 2015), and introduced into seed-derived calli of *japonica* rice cultivar Nipponbare by the method previously reported (Hiei et al. 1994). All plants were grown in growth chambers at 30°C day and 25°C night temperature and 70% relative humidity with a daylength of 12 hours.

### Pollen and seed fertility tests

For pollen fertility, anthers extracted from fixed panicles with Carnoy’s fluid were squashed in I_2_-KI solution, and viable pollen grains stained were counted under the light microscope BX50 (Olympus). For seed fertility, the ratio of fertile spikelet numbers was counted in each of five panicles and averaged.

### Cytological observations

For observations of chromosome behaviours and callose deposition on anther sections, anthers were fixed in 4% paraformaldehyde (PFA)/1xPBS. After removal of lemma and palea of florets, anthers were dehydrated in ethanol graded series for 30-60 min each, followed by infiltration in embedder with hardener1 of Technovit 7100 (Kulzer Technique), overnight on rotor at 4°C. The solution was replaced with fresh Technovit with hardener1 every 6 hours and incubated overnight with on rotor at 4°C. Then, anthers were transferred to a cryo-dish with Technovit polymerised by addition of harderner2 and placed at 50-60°C for hardening. Plastic embedded sections with 4-6μm thickness were taken using the microtome R2255 (Leica), and air dried at room temperature. The section was stained for 25-30 min with 0.01% aniline blue (Sigma Aldrich) in 0.1M K_3_PO_4_ (pH 12) for callose, or with a drop of 1.5μg/mL 4’,6-diamidino-2-phenylindole (DAPI)/Vector shield (Vectorlabs) for chromosome observation. The images were captured under the confocal laser scanning microscope system FV300 (Olympus), and processed with ImageJ (https://imagej.nih.gov/ij/docs/intro.html).

For chromosomal spreads, whole panicles were fixed in Carnoy’s fluid and stored at 4°C until use. Fixed anthers incubated with 0.1% FeCl_2_ overnight were squashed in acetocarmine solution (1% (w/v) carmine (Merk)/45% acetic acid) with forceps on a clean glass slide. After quick removal of anther-wall remnants, a suspension of released PMCs was covered with a cover slip, and gently heat-treated followed by gentle thumb compression. The images of chromosomal spreads were captured under the light microscope BX50 with DP2-SAL CCD camera system (Olympus). The number of PMCs each classified by certain phenotypes in chromosome behaviours was counted and used for quantification together with those observed in plastic-embedded sections.

### RT-qPCR

To quantify the transcript levels of rice genes, anthers and tissues were collected, immediately frozen in liquid nitrogen in a 2mL tube with 2mm beads, and homogenized on automated shaker BMS-A20TP (Bio Medical Science) at 1100rpm for 2min. RNAs were extracted by TRIZOL RNA extraction kit method according to manufacturer’s instruction (Invitrogen) and supplied to Super Script III First-Strand synthesis system (Invitrogen) for cDNA library construction. RT-qPCR was performed using Real Time System TP800 (TAKARA bio systems), according to manufacturer’s instruction. All primer sequences used for RT-qPCR were shown in Table S2. *Rice Actin1* (*RAc1*) gene was used as an internal control to normalize the expression levels at all meiosis stages quantified.

### Antibody production

To produce an antibody specific for OsGSL5 of 1910 amino acids (aa), the cDNA sequence encoding 1009-1260 aa position was amplified using the above cDNA library as a template (Fig. S4, Table S2), and cloned into pDEST17 vectors (Invitrogen). The His-tagged protein expressed in *E. coli* strain BL21-AI (Invitrogen) was purified using Ni-NTA agarose resin (FUJIFILM), and immunised to rabbits and guinea pig.

To observe telomere behaviours in PMCs, the antibody was raised against rice PROTECTION OF TELOMERE1 (OsPOT1), which is encoded by a single gene locus *Os04g0467800* (*LOC_Os04g39280*.*1*), while Arabidopsis genome has two paralogous loci, *POT1a* and *POT1b* (Shakirov et al. 2005). Procedures to raise antisera are same as above.

### Immunofluorescent staining of PMCs and tissue sections

Young rice panicles were fixed with 4%PFA/1xPMEG (25mM PIPES, 5mM EGTA, 2.5mM MgSO4, 4% glycerol, and 0.2% DMSO, pH 6.8), followed by washing 6 times with 1xPMEG, and stored at 4°C until use (Nonomura et al. 2006). For immunofluorescence of PMCs, fixed anthers were treated with the enzyme cocktail of 2% cellulase Onozuka-RS (Yakult)/0.3% pectolyase Y-23/0.5% macerozyme-R10 (FUJIFILM Wako)/0.375% Cytohelicase (Sigma-Aldrich) in 1xPME (same with 1xPMEG except for excluding glycerol) for permeabilization on MAS-coating glass slide MAS-02 (Matsunami Glass), squashed by forceps and used for immunostaining, as described in Nonomura et al. (2006).

Immunostaining of anther sections was done as described by Tsuda and Chuck (2019) with minor modifications. Briefly, anthers were dehydrated in ethanol graded series and Histo-Clear (Cosmo bio co. Ltd), embedded into Paraplast paraffin wax (McCormick Scientific). Paraffin blocks containing anther samples were trimmed and stored at 4°C until use. The blocks were sectioned into 8-10μm thickness by the microtome. Dewaxed and rehydrated samples were incubated with primary antibodies. The rabbit anti-OsGSL5, the rabbit anti-OsPOT1 (this study), the rabbit anti-PAIR2 (Nonomura et al. 2006) and rat anti-ZEP1 antibodies (Nonomura et al. 2011) were diluted to 1/100, 1/3000, 1/3000 and 1/1000, respectively, with 3%BSA/1xPMEG and used as primary antibodies. For callose immunostaining, monoclonal antibody specific to β-1,3-glucan (callose) (Biosupplies Australia) was diluted to 1/1000 and used as a primary antibody was used.

In both squash and sectioning methods, secondary antibodies Alexa fluor 488 (Abcam) and Cy3-conjugated IgG (Merck) of 1/200 dilution was used for detection. Immunofluorescent images were captured by Fluoview FV300 CSLM system (Olympus) and processed with ImageJ.

## Acknowledgments

We thank Dr. Norio Komeda (NIG, SOKENDAI) for helping the production of anti-OsPOT1 antisera, and Dr. Yoshihisa Oda (NIG) for reading manuscript and giving useful comments. The *mel2* mutant used in this study was provided by NIG with support from National BioResource Project (NBRP) Rice, AMED, Japan. This work was partly supported by JSPS KAKENHI Grant No. 21H04729 and 18H02181 (to K.I.N), and Bilateral Programs Grant No. JPJSBP120213510 (to K.I.N). We also thank MEXT (The Ministry of Education, Culture, Sports, Science and Technology, Govt. of Japan) for the support.

## Author contributions

H.S, M.M and K.I.N have designed the research work. H.S carried out most of the experiments. K.T has helped in the production of antibody against GSL5. M.M has guided in immunostaining experiments. H.S and K.I.N wrote the manuscript. M.M and K.T helped in drafting the manuscript.

## Data Availability

The data presented in this study are available from the corresponding author upon reasonable request.

## Figure legends

**Supporting figure 1. Transcript levels of *OsGSL5* gene quantified by RT-qPCR in WT and *mel2* anthers**.

**Supporting figure 2. Callose accumulation in WT and *mel*2 anthers**.

**Supporting figure 3. Expression profile of *OsGSL5* gene at various tissue developmental stages in WT rice plants**.

**Supporting figure 4. OsGSL5 protein structure**.

**Supporting figure 5. Pollen viability in WT and *osgsl5* mutant**.

**Supporting figure 6. The expression level of *OsGSL5* transcripts in various vegetative tissues including male and female organs in WT and *osgsl5-2* plants**.

**Supporting figure 7. Anther development is unaffected in *osgsl5* mutants**.

**Supporting figure 8. Callose deposition at its beginning stage during premeiotic interphase in WT anthers**.

**Supporting figure 9. Immunostaining of OsGSL5 protein**.

**Supporting figure 10. Aberrant chromosome behaviors detected in *osgsl5* mutant anthers by chromosome spreading technique**.

**Supporting figure 11. Transcript levels of various meiotic genes in WT and *osgsl5-2* anthers**.

**Supporting figure 12. Telomere bouquet structures visualized at late prophase stage**.

**Supporting table 1. Oligo DNA sequences for guide RNA constructions and PCR primers employed in this study**.

**Supporting table 2. Seed fertility of WT, *osgsl5-2* and *osgsl5-3* plants**.

**Supporting table 3. PMC counting data on the basis of Figure 6 graphs**.

